# From stress to depression: Development of extracellular matrix-dependent cognitive impairment following social stress

**DOI:** 10.1101/806935

**Authors:** Maija-Kreetta Koskinen, Yvar van Mourik, August Benjamin Smit, Danai Riga, Sabine Spijker

## Abstract

Stress can predispose to depressive episodes, yet the molecular mechanisms regulating the transition from the initial stress response to a persistent pathological depressive state remain poorly understood. To shed light on this stress-to-depression transition process, we profiled the development of an enduring depressive-like state in rat by assessing affective behavior and hippocampal function during the 2 months following social defeat stress. In addition, we measured remodeling of hippocampal extracellular matrix (ECM) during this period, as we recently identified ECM changes to mediate cognitive impairment during a sustained depressive-like state. We found affective disturbance and cognitive impairment to develop disparately after social stress. While affective deficits emerged gradually, spatial memory impairment was present both early after stress and during the late-emerging chronic depressive-like state. Surprisingly, these phases were separated by a period of normalized hippocampal function. Similarly, the SDPS paradigm induced a biphasic regulation of the hippocampal ECM coinciding with hippocampus-dependent memory deficits. Early after stress, synaptic ECM proteins and the number of perineuronal nets enwrapping parvalbumin-expressing interneurons were decreased. This was followed by a recovery period without ECM dysregulation, before subsequent decreased metalloproteinase activity and ECM build-up, previously shown to impair memory. This suggests that intact hippocampal function requires unaltered ECM levels. Together our data 1) reveal a dichotomy between affective and cognitive impairments similar to that observed in patients, 2) indicate different molecular processes taking place during early stress and the chronic depressive-like state, and 3) support a role of the ECM in mediating long-lasting memory-effects of social stress.

## Introduction

Major depressive disorder (MDD) is a highly prevalent and debilitating neuropsychiatric disorder with a complex set of symptoms and high rates of relapse^1^. MDD is characterized by persistent disturbances in the affective domain, including depressed mood and anhedonia^1^. In addition to these well-characterized affective symptoms, cognitive dysfunction is prominent among depressed patients, affecting multiple domains, such as executive function, attention, memory and learning^2, 3^. Despite the high occurrence and disabling effects on everyday life of patients, cognitive dysfunction in depression has received little attention and therapeutic strategies against depression are mainly aimed at alleviating its mood-related symptoms. This poses serious limitations as residual cognitive symptoms often persist after improvement of mood symptoms and prevent functional recovery of patients^4, 5^. Moreover, these residual symptoms are associated with an increased risk of relapse and recurrence of depressive episodes^6^. Taken together, increasing data suggest that rather than being an epiphenomenon of affective symptoms, cognitive dysfunction represents a core trait of the disease that is necessary to tackle in order to reach full recovery and prevent relapse.

Although the etiology of MDD remains elusive, stressful life experiences have been strongly linked to the development of depression and other neuropsychiatric disorders^7^. The hippocampus is highly susceptible to the effects of stress and given its critical involvement in cognitive function (e.g., spatial and temporal order processing), stress-induced hippocampal abnormalities appear to have a strong contribution to the cognitive deficits associated with MDD^8, 9^. For example, attenuated hippocampal activity during hippocampus-dependent tasks has been demonstrated in MDD patients^10, 11^, underscoring the potentially causal relation between deficits in recollection and declarative memory and hippocampal atrophy. Similarly, in chronically stressed animals, hippocampal memory deficits are present alongside structural changes and altered activity of hippocampal neurons^12^. Despite the vast data describing these stress-induced depression-related changes, the neural substrates and mechanisms that underlie the transition from initial stress exposure to a pathological depressive state remain elusive. In particular, the molecular mechanisms that propel these enduring effects of stress are poorly characterized as preclinical studies have largely focused on the short-lasting effects of acute and/or chronic stress.

Recently, we employed the Social Defeat-induced Persistent Stress (SDPS) model in rats to investigate the molecular mechanisms that underlie hippocampus-dependent cognitive dysfunction during a sustained depressive-like state^13^. In this preclinical model, brief social defeat stress is combined with a prolonged social isolation period that together result in enduring affective deficits and cognitive dysfunction, thereby recapitulating the core depression symptoms. Importantly, these deficits are prominent months after the last social defeat stress exposure, reminiscent of the human depressive state that can emerge and persist long after a stressful period^14, 15^. As a novel pathological mechanism, we identified changes in hippocampal extracellular matrix (ECM), which mediate these cognitive impairments during the sustained depressive-like state^13^.

The brain ECM is a heterogeneous molecular network that forms a scaffold around neurons and synapses, supporting structural stability and regulating plasticity^16, 17^. The pivotal role for ECM in mediating experience-dependent plasticity in the adult brain has become increasingly apparent. In particular, perineuronal nets (PNNs), lattice-like matrix structures predominantly coating parvalbumin-expressing interneurons^18^, have been shown to gate the closure of critical periods^19^, to sustain fear memories^20^ and to regulate addiction-related memories^21^. Furthermore, increasing evidence supports the involvement of abnormal ECM in diseases with disturbed cognitive processing, as diverse as schizophrenia^22^ and Alzheimer’s disease^23^.

In the present study, we studied the development of SDPS-induced depressive-like state by assessing affective behavior, as well as cognitive function during the days and weeks following social defeat stress. In addition to this temporal profiling of the core depressive-like symptoms, we characterized ECM remodeling over this same timeframe in order to understand how aberrant ECM that underlies the cognitive deficit at the sustained depressive-like state, develops following initial social defeat stress.

## Materials and Methods

The full method section can be found in the Supplementary Materials.

### Animals and the social defeat-induced persistent stress (SDPS) paradigm

All experiments were approved by the central ethics committee of the Netherlands and the Animal Users Committee of the Vrije Universiteit Amsterdam, in accordance with the relevant guidelines and regulations. The SDPS paradigm was carried out as previously described^13^, where male male Wistar rats (≥ 9 weeks; Envigo, Netherlands) were used in a resident-intruder paradigm with male Long-Evans rats (>4 months; Charles River, UK) as residents. The SDPS rats underwent 15-minute social defeat sessions daily with physical contact for 5 minutes. The defeat was repeated for five consecutive days and each day a new resident was used. From the first defeat session onwards, the SDPS rats were single-housed until the end of the experiment.

### Behavioral testing

All behavioral testing was performed during the dark phase of a 12 h light-dark cycle (lights on at 7 PM), under a dim red light as described previously^13^. Affective function was tested with the Social Approach Avoidance (SAA test), in which exploration and approach to a caged unfamiliar Long-Evans rat was measured during the first minute of the test as the time spent near the social target *vs*. time spent near the empty box; interaction ratio: social target zone / (social target zone + empty box zone). Cognitive function in terms of spatial memory was assessed with the Object Place Recognition (OPR test), in which exploration and approach of a relocated object was analyzed during the first minute of the test as time spent exploring the relocated object compared to the stable object; Discrimination index: relocated object / (relocated object + stable object).

### Immunohistochemistry

Following transcardial perfusion with ice-cold 4% PFA in PBS and and overnight post-fixation, brains were transferred to 30% sucrose in PBS at 4 °C until sectioned and processed for immunohistochemistry as described previously^13^. Free-floating sections were incubated with primary antibodies (mouse anti-chondroitin sulfate proteoglycan 1:1,000, cat-301 MAB5284; rabbit anti-parvalbumin 1:1,000, Swant #235) overnight at 4 °C. After washing, sections were incubated with fluorescent-conjugated secondary antibodies (anti-mouse-Alexa-488 1:400, Invitrogen A11001; anti-rabbit-Alexa-568 1:400, Invitrogen A11011) for 2 h at RT. Thereafter, the sections were washed 4 times 10 minutes with PBS at RT and mounted.

Images were acquired on a fluorescent microscope (Leica DM5000), and analyzed by Fiji software^24^, using automated threshold and particle analysis to detect the number of PNN^+^ and PV^+^ cells and their double immunoreactivity. False-positive cells were excluded manually during the analysis. During image acquisition and cell quantification, the researcher was blind to the experimental groups.

### Tissue preparation, immunoblotting, MMP extraction & zymography

Dorsal hippocampus^25^ samples were homogenized to either isolate synaptosomes for immunoblotting using a sucrose gradient as described previously^13^ or to isolate ECM-bound MMPs for zymography as described^26^.

#### Immunoblotting

Protein concentration was determined (Bradford protein assay) and 10 µg of protein was loaded per sample for electrophoresis. The following primary antibodies were used: rabbit anti-aggrecan (1:700, AB1013, Abcam); guinea-pig anti-brevican (1:2,000; generously provided by C.I. Seidenbecher, Magdeburg), mouse anti-neurocan (1:1,000, N0913 Sigma); mouse anti-phosphacan (1:1,000; 3F8, Developmental Studies Hybridoma Bank); mouse anti-versican (1:1,000; 75-324, NeuroMab); mouse anti-Tenascin-R (1:2,000, mTNR-2 Acris Antibodies); rabbit anti-Hapln1 (1:1,000, ab98038 Abcam). After incubation with horseradish peroxidase-conjugated secondary antibody (1:10,000; Dako, Glostrup, Denmark) and visualization with Femto Chemiluminescent Substrate (Thermo Scientific, Rockford, IL, USA), blots were scanned using the Li-Cor Odyssey Fc (Westburg, Leusden, The Netherlands) and analyzed with Image Studio (Li-Cor, Lincoln, NE, USA). Total protein was visualized using trichloro-ethanol staining and scanned using a Gel Doc EZ imager (BioRad, Herculus, CA, USA) and analyzed with Image Lab (BioRad) to correct for differences in sample loading. Data were subsequently log_2_-transformed and presented *vs*. those of control samples. In addition, data were compared to the log_2_-transformed quantification of perisynaptic ECM protein levels, as shown previously^13^.

#### MMP extraction & zymography

Tissue was homogenized (50 mM Tris pH 7.4, 10 mM CaCl_2_ and 0.25% Triton X-100) and centrifuged (6,000x *g*, 30 minutes at 4 °C). The pellet was resuspended (50 mM Tris, pH 7.4, and 0.1 mM CaCl_2_), heated (60 °C, 15 minutes) and centrifuged (10,000x *g*, 30 minutes at 4 °C). After this, the supernatant was recovered and the protein concentration was determined using Bradford. Thereafter, proteins were precipitated (60% ethanol, 1 minute at 4 °C) and centrifuged (15,000x *g*, 5 minutes at 4 °C). Finally, the pellet was solubilized in non-reducing sample buffer (2% SDS) and heated (37 °C, 15 minutes) before loading on an SDS-PAGE gel containing gelatin (0.1% gelatin, 8% SDS); samples (10 µg of protein) and recombinant mouse MMP-9 as positive control (5 ng, ab39309, Abcam). The gels were washed with 2.5% Triton X-100 (2x 20 minutes) and then incubated for 7 days (50 mM Tris, pH 7.5, 10 mM CaCl_2_, 1 μM ZnCl_2_, 1% Triton X-100, and 0.02% sodium azide; 37 °C, 80 rpm). Incubation was followed by Coomassie staining and destaining (5% HAc) until clear bands were visible. Gels were scanned and analyzed using Gel Doc EZ imager (Biorad, Herculus, CA, USA), and normalized based on Coomassie input.

### Statistics

All data were analyzed using IBM SPSS Statistics 24. For group comparisons, two-tailed Student’s t-tests (with or without correction for unequal variation) were applied for normally distributed data and Mann-Whitney U-tests otherwise. Data were checked for normality using the Saphiro-Wilk test. All group data are depicted as mean±SEM, with individual data on top. Statistical significance level was set for *P*-values<0.05, trend for 0.05<*P*-value<0.10. Correlations were made for SAA and OPR behavior per batch of animals, using Spearman rank (n<15), or Pearson’s product moment (n≥15). Details of all statistical testing can be found in Supplementary Table S1, and individual data points are given in Supplementary Table S2.

## Results

### Affective dysfunction and cognitive impairment develop disparately following social defeat stress

To characterize the development of depressive-like symptoms following a stressful experience, we performed a temporal analysis and assessed affective behavior and cognitive function at several time-points (24 h / 48 h, 2, 4 and 8 weeks) following the last social defeat encounter (**Fig. 1a**). The effect of SDPS on affective behavior was tested by the social approach avoidance (SAA) test, where interaction towards an unfamiliar Long-Evans rat is used to measure motivation for social interaction. Cognitive function was assessed by the object place recognition (OPR) test with a 15-minute retention interval. This test measures short-term spatial memory and is heavily dependent on hippocampal function. In order to avoid carryover effects from repeated testing, independent groups of animals were used for each time-point. In addition, these low-stress tests allowed analysis of ECM expression in absence of stress-by-test-induced effects.

**Figure 1.**
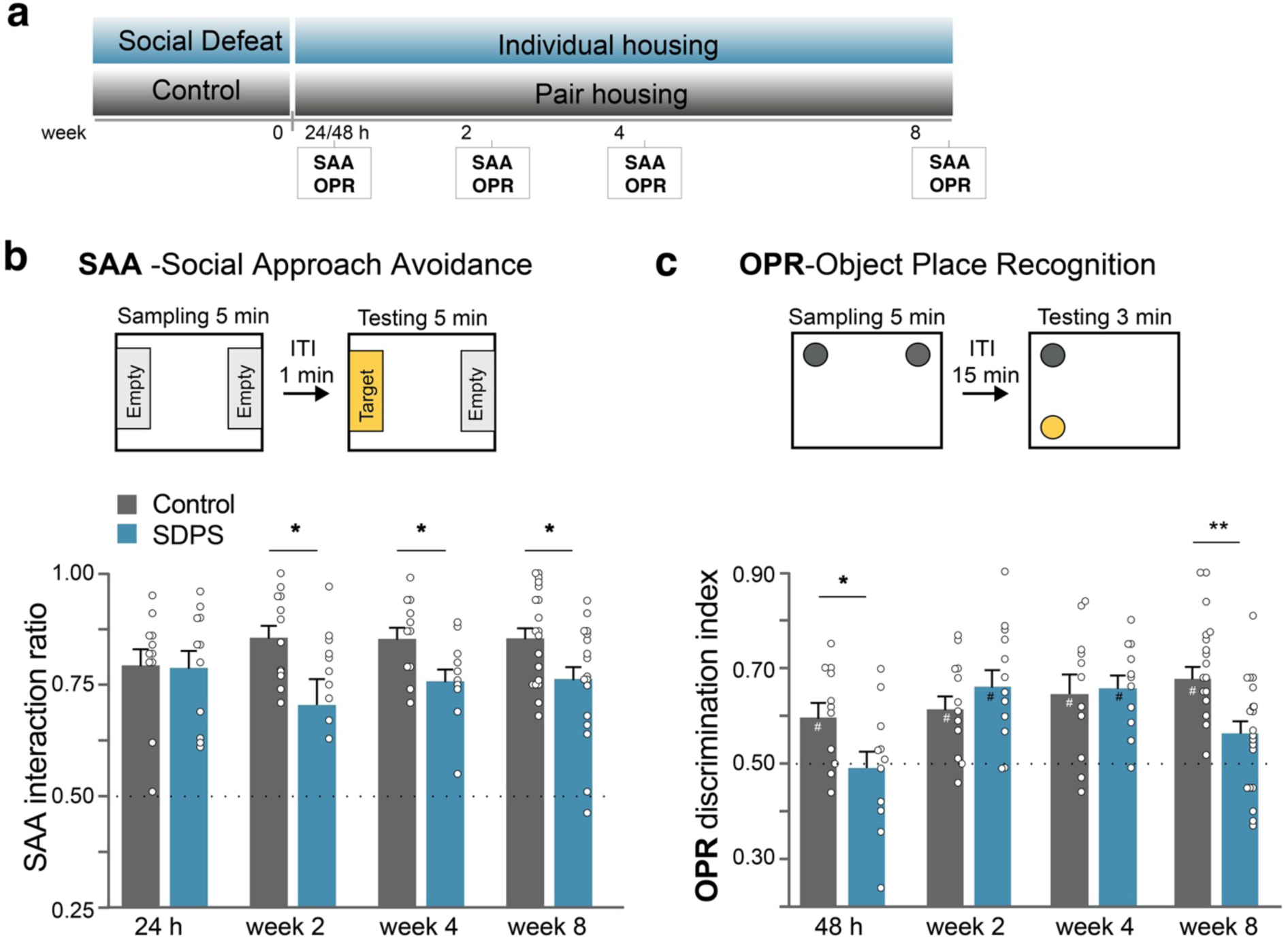
Affective deficit and cognitive impairment develop disparately after social defeat stress. **a)** Adult rats were subjected to a 5-day social defeat paradigm followed by an 8-week period of individual housing. Control rats were not exposed to social defeat and remained in pair housing throughout the experiment. Behavior was assessed with Social Approach Avoidance (SAA) and Object Place Recognition (OPR) tests at four different time-points: 24 h / 48 h, 2 weeks, 4 weeks and 8 weeks after the last social defeat session. **b)** Affective deficits emerged gradually following SDPS (Social Defeat-induced Persistent Stress). At 24 h after the last social defeat session, no difference in approach towards a social target between SDPS and control group was present. At week 2 post-defeat and thereafter decreased approach towards a social target was observed in the SDPS group **c)** SDPS impaired spatial memory in a temporally dynamic manner. The SDPS rats failed to discriminate between a stable and relocated object both shortly (48 h) after social defeat stress and at the chronic depressive state (week 8), whereas at week 2 and 4 post-defeat the SDPS rats showed a preference for the relocated object similar to the control group. Behavior at each time-point was assessed in an independent group of SDPS and control rats. Student’s t-test or Mann-Whitney U test were used to test statistical significance between SDPS and control group at each time-point. #P<0.05 compared to a fictive group with a mean of 0.5 and equal variation; *P<0.05; **P<0.01. Data are expressed as mean±SEM.

A day after (24 h) the last social defeat exposure, control and SDPS animals showed similar interaction time spent in the SAA test (control *vs*. SDPS, *P*=0.935), indicating that recent stress does not affect approach behavior towards an unfamiliar social target (**Fig. 1b**). At 2 weeks post-defeat, the SDPS group displayed decreased time spent with the social target (control *vs*. SDPS, *P*=0.028), demonstrating the emergence of diminished motivation for social interaction and exploration. Similarly, at week 4 and 8 post-defeat, the SDPS rats showed decreased interaction ratios (control *vs*. SDPS, week 4 *P*=0.023; week 8 *P*=0.026). Taken together, these results demonstrate a gradually developing, yet persistent, effect of SDPS on affective behavior as observed previously^13, 27^.

The next day (48 h after the last social defeat session), the SDPS group showed a deficit in the OPR test, assessing short-term spatial memory. A significant group effect was found (control *vs*. SDPS, *P*=0.041), as the SDPS rats spent significantly less time exploring the relocated object when compared with the control group, thereby demonstrating an impairment in hippocampus-dependent spatial memory. Furthermore, whereas the control group showed a preference for the relocated object (control, *P*=0.029 *vs*. a fictive control with a mean of 0.5 and equal variation^28^), the SDPS group failed to perform this task by showing an OPR discrimination index around chance levels (SDPS, *P*=0.908 *vs*. a fictive control) (**Fig. 1c**). Surprisingly, at week 2 post-defeat, the SDPS group displayed normal short-term memory retention (SDPS, *P*=0.003 *vs*. a fictive control) similar to the control group (control, *P*=0.008 *vs*. a fictive control), while no between group differences were found (control *vs*. SDPS, *P*=0.298). Also, at week 4 post-defeat both groups showed a preference for the relocated object (control, *P*=0.018 *vs*. a fictive control; SDPS, *P*<0.001 *vs*. a fictive control; and control *vs*. SDPS, *P*=0.789). In accordance with our previous study^13^, 8 weeks after the last social defeat stress encounter, SDPS rats showed impaired memory function as indicated by decreased exploration of the relocated object compared with the control group (control *vs*. SDPS, *P*=0.002) and a loss of preference for the displaced object (SDPS, *P*=0.067 *vs*. a fictive control), whereas the control group retained information over the object’s location (control, *P*<0.001 *vs*. a fictive control).

Taken together, the current data show that hippocampal function is restored for a limited period of time after the short-term stress effects subside and before the long-term depression-related changes emanate. No significant correlation between SAA and OPR behavior was observed in any of the time-points studied (Supplementary Fig. S1), indicating a degree of independence between these two behavioral domains, as shown previously^27^.

### Cognitive impairment coincides with dysregulation of hippocampal extracellular matrix

Our previous study identified increased levels of hippocampal ECM at the depressive-like state evident months after the last social defeat exposure^13^. Specifically, SDPS induced an increase in the number of PNN-coated PV-expressing interneurons in the dorsal CA1 hippocampal subregion, together with an increase in the expression of ECM proteins in the synaptic membrane fraction of the dorsal hippocampus. Moreover, we showed that enzymatic disruption of CA1-ECM could restore the number of PNNs and, importantly, rescue the SDPS-induced hippocampal memory deficits, thereby demonstrating a causal relationship between hippocampal ECM dysregulation and cognitive dysfunction during the sustained depressive-like state. However, at what moment after the defeat stress this ECM increase emerged was not clear.

To unravel the progression of SDPS-induced dysregulation of hippocampal ECM, in the present study we profiled temporal changes in the ECM at 72 h, 2, 4 and 8 weeks after social defeat stress. We quantified the number of the PNN^+^/PV^+^-cells in the dorsal CA1 region (**Fig. 2a**). Early after social defeat stress (72 h), a reduced number of PNN^+^/PV^+^-cells was found in the SDPS animals compared with the control group (control *vs*. SDPS, *P*=0.030). This shows that social defeat stress induces an immediate ECM remodeling resulting in decreased number of PNN^+^/PV^+^-cells. At week 2 and 4 post-social defeat, no difference in the number of PNN^+^/PV^+^-cells between the control and SDPS groups was present (week 2, control *vs*. SDPS, *P*=0.941; week 4, control *vs*. SDPS, *P*=0.196), revealing a recovery of the perisomatic ECM after the cessation of acute stress. Moreover, as shown previously^13^, an increased number of PNN^+^/PV^+^-cells was present 8 weeks following stress, characterizing the sustained depressive-like state (week 8, control *vs*. SDPS, *P*=0.021) (**Fig. 2b,c**). The density of PV^+^ interneurons was not affected in any of the time points studied (72 h, control *vs*. SDPS, *P*=0.977; week 2 control *vs*. SDPS, *P*=0.155; week 4 control *vs*. SDPS, *P*=0.110; week 8 control *vs*. SDPS, *P*=0.333; **Fig. 2d**), indicating a specific regulation of PNNs by SDPS both early after stress and at the phase when the depressive-like state appears (*cf*. Fig. 1).

**Figure 2.**
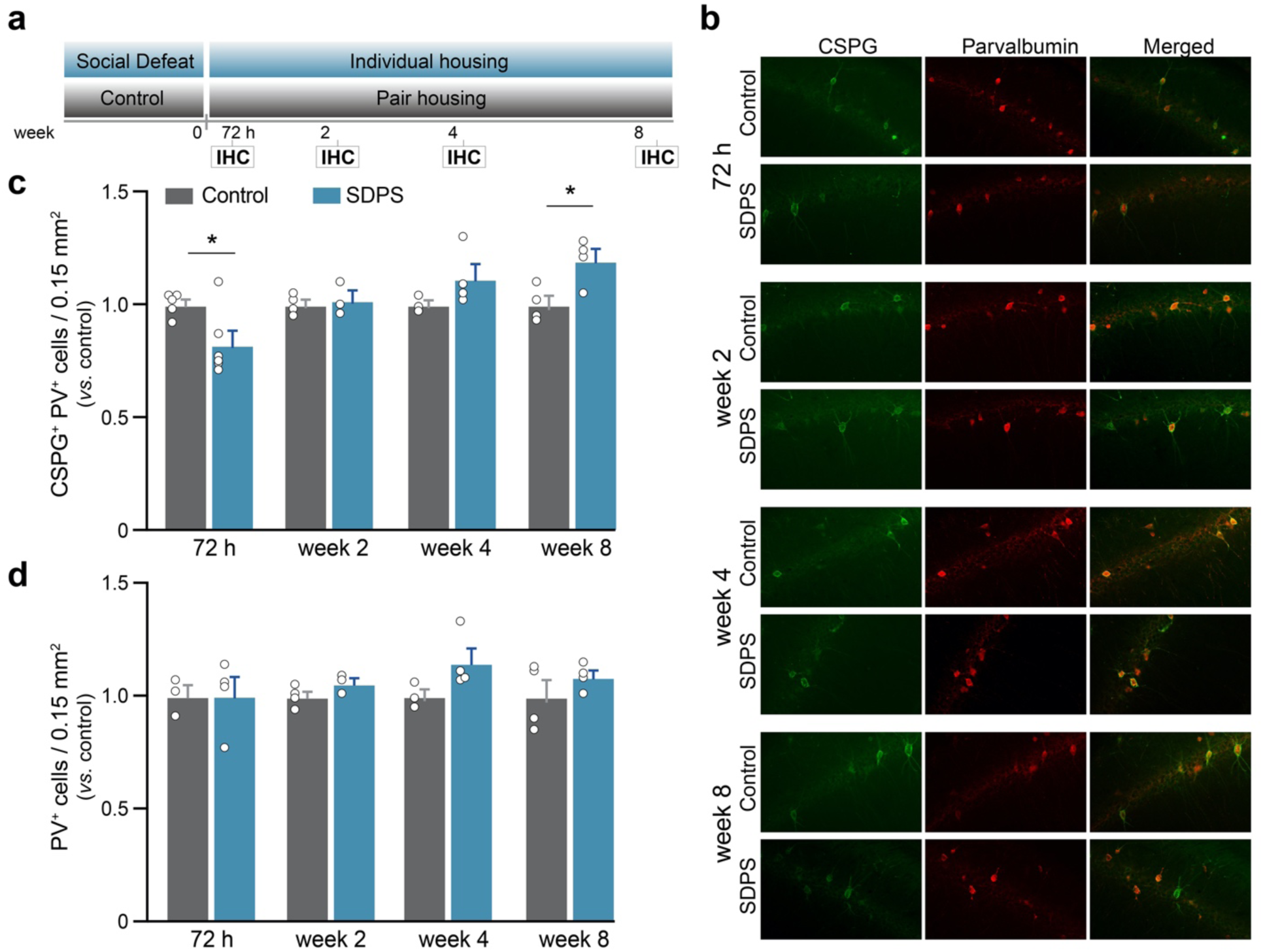
SDPS induces a temporally dynamic regulation of CA1 perineuronal nets. **a)** The number of chondroitin sulfate proteoglycan (CSPG^+^)-rich perineuronal nets (PNNs) and parvalbumin-expressing (PV^+^) interneurons were quantified with immunohistochemical (IHC) analysis at four time-points: 72 h, 2 weeks, 4 weeks and 8 weeks after the last social defeat session. **b)** Decreased number of CSPG^+^PV^+^ -cells was present at 72 h after stress in the SDPS group. At week 2 and 4 no difference in the number CSPG^+^PV^+^-cells was observed. At the chronic depressive state (week 8) an increased number of CSPG^+^PV^+^-cells was found in the SDPS group, revealing a biphasic regulation of PNNs in response to SDPS. **c)** SDPS did not alter the total number of PV-expressing cells in any of the time-points studied. Student’s t-test was used to test statistical significance between SDPS and control group at each time-point. *P<0.05. Data are expressed as mean±SEM.

Apart from the perisomatic PNNs surrounding PV-interneurons, perisynaptic extracellular space is enriched with ECM proteins that are vital for synaptic function^29, 16^. Therefore, we also investigated the expression of ECM proteins in the synaptic dorsal hippocampus fraction 72 h and 2 weeks following social defeat stress (**Fig. 3a**). We found the expression of the perisynaptic ECM to follow an identical pattern with that of PNNs. Namely, early after stress we found decreased expression of ECM proteins, followed by a transient recovery period during which perisynaptic ECM regulation was absent, before increasing at the long-term (**Fig. 3b,d**). Specifically, at the early time-point (72 h), synaptic expression of several chondroitin sulfate proteoglycans (CSPG) from the lectican family, including brevican (Bcan), neurocan (Ncan) and phosphacan (Phcan) was reduced in the SDPS group (*vs*. control: Bcan, *P*=0.039; Ncan, *P*=0.029; Phcan, *P*=0.022; Acan, *P*=0.450). Similarly, the expression of the link-proteins Tenascin-R (TnR) and Hyaluronan and proteoglycan link protein 1 (Hapln1) was decreased shortly after social stress in the SDPS group (*vs*. control: TnR_160_, *P*=0.007; TnR_180_, *P*=0.047; Hapln1, *P*=0.021) (**Fig. 3c,d**). At week 2 post-defeat, no difference in the expression of synaptic ECM proteins between the control and SDPS groups was present (*vs*. control: Bcan, *P*=0.993; Ncan *P*=0.760; Phcan, *P*=0.671; Acan, *P*=0.300; TnR_160_, *P*=0.327; TnR_180_, *P*=0.629; Hapln1, *P=*0.491), illustrating a transient recovery of perisynaptic ECM protein levels, similar to what was observed for the number of PNNs.

**Figure 3.**
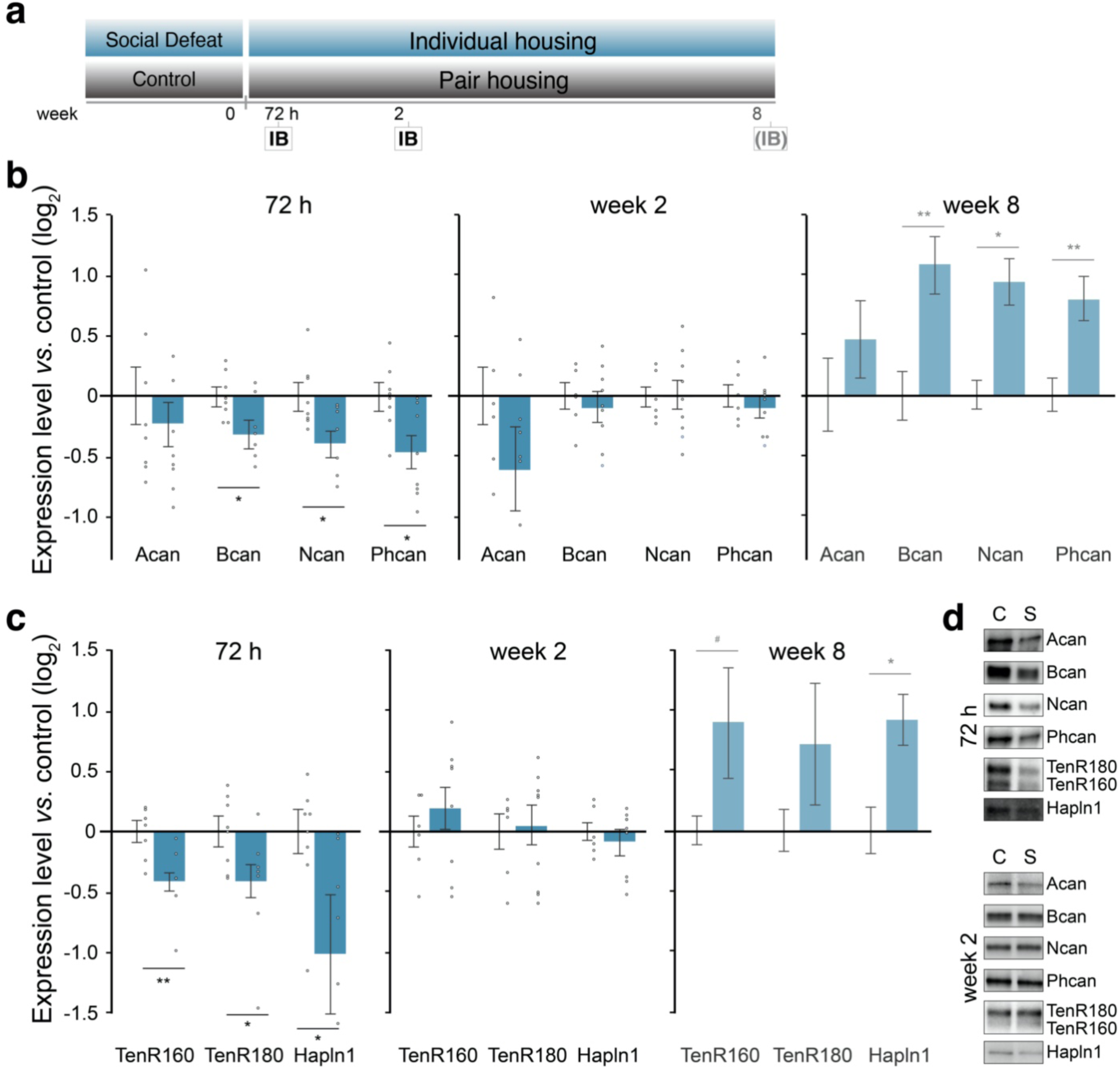
SDPS induces a biphasic regulation of perisynaptic extracellular matrix. **a)** Immunoblot (IB) analysis was performed at two time-points: 72 h and 2 weeks after the last social defeat session, and was compared to the log_2_-transformed quantification of perisynaptic ECM protein levels 8 weeks post-defeat^13^. **b-c)** Decreased expression of chondroitin sulfate proteoglycans Brevican (Bcan), Neurocan (Ncan) and Phosphacan (Phcan), together with glycoproteins Tenascin (TnR) and Hyaluronan and proteoglycan link protein 1 (Hapln1) were present shortly after stress (72 h) in the SDPS group. At week 2 no difference in protein expression level was observed. **d)** Example blots of each protein at the 72 h and 8 week post-defeat time-point for Control (C) and SDPS (S). Full blots, as well as corresponding gels for total protein (sample normalization) are available in Supplementary Figures 2 and 3. Student’s t-test was used to test statistical significance between SDPS and control group at each time-point. * P<0.05; **P<0.01. Data are expressed as mean±SEM.

Taken together, this temporal profile of hippocampal ECM organization shows that social defeat stress induces a persistent, yet dynamic ECM remodeling both at the perisynaptic, as well as at the pericellular level.

### MMP-2 activity is decreased during the sustained depressive-like state

Matrix metalloproteinase-mediated breakdown of ECM assemblies is an essential mechanism by which ECM remodeling is maintained^30, 31^. To assess whether altered MMP activity potentially drives the SDPS-associated ECM changes, we measured gelatinolytic activity of MMP-2 and MMP-9 using in-gel zymography at the two time-points showing major ECM dysregulation (**Fig. 4a**). Early after social stress, we did not find a significant difference in the activity of either MMPs (*vs*. control: MMP-2, 118%, *P*=0.619; MMP-9, 139%, *P*=0.301; **Fig. 4b,d**). Notably, at 8 weeks after social defeat, we found decreased MMP-2 activity in the SDPS animals compared with controls (*vs*. control: 76%, *P*=0.043), whereas no difference in MMP-9 activity was present (*vs*. control: 84%, *P*=0.239; (**Fig. 4c,d**). Together, our data suggest that increased build-up of the perisynaptic and pericellular ECM, as observed during the sustained depressive-like state, could result from decreased MMP-2-mediated breakdown of the ECM in the hippocampus, whereas the early ECM effects after social defeat are possibly regulated by an MMP-2/MMP-9 independent mechanism.

**Figure 4.**
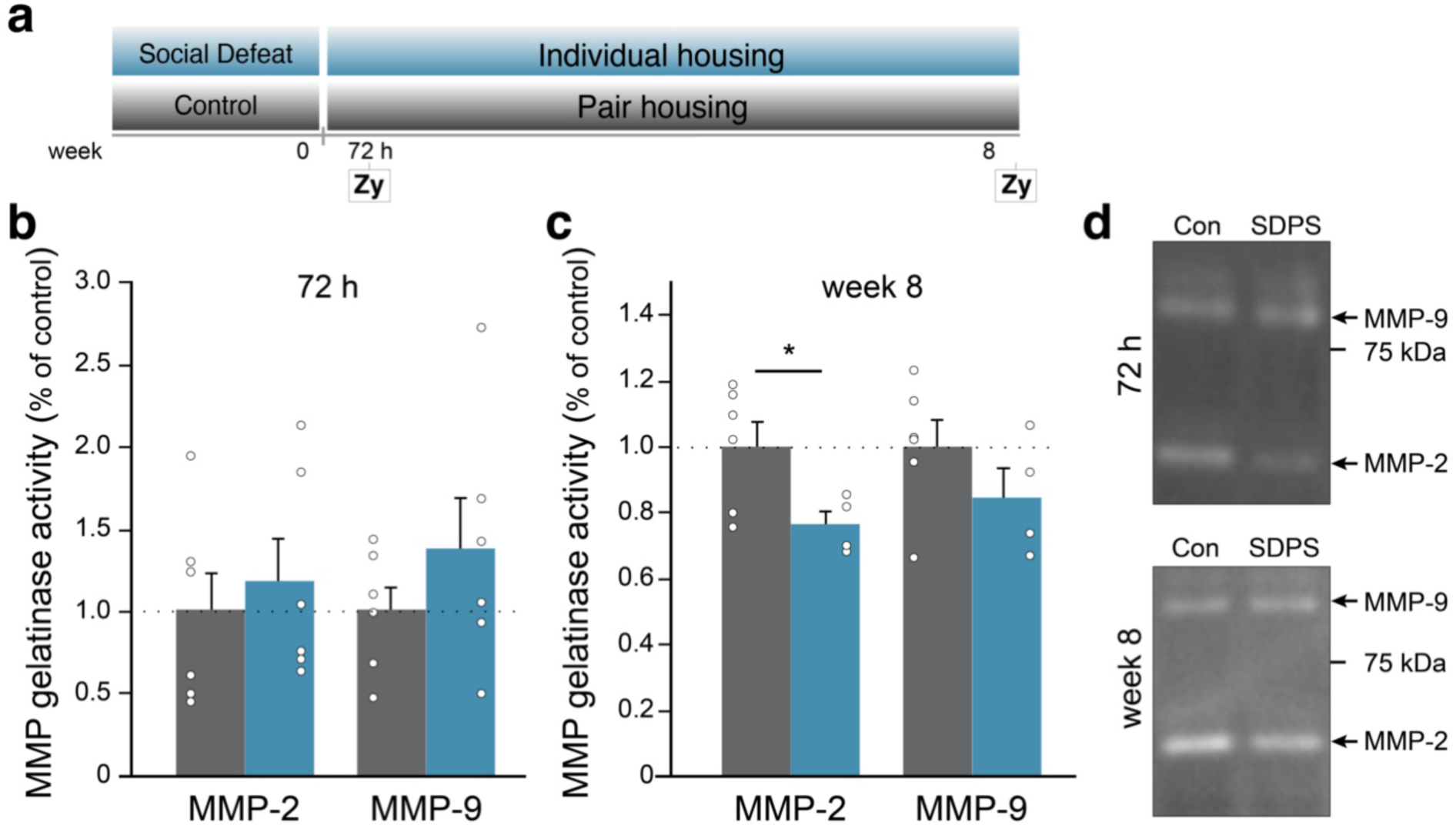
MMP-2 activity is reduced during the sustained depressive-like state. **a)** In-gel zymography assays (Zy) were performed at 72 h and 8 weeks after social defeat. **b)** No difference in MMP-2 or MMP-9 activity was present shortly after social defeat. **c**) At week 8 post-defeat, decreased MMP-2 activity was found in the SDPS group, without MMP-9 regulation. * P<0.05. Data are expressed as mean±SEM. **d**) Examples with equal loading showing the mature form of MMP-2 and MMP-9 (white bands), and the position of the 75 kDa marker band for each time point. Note that for the 72 h time-point a large variation between samples was observed. Full gels are available in Supplementary Figure 4.

## Discussion

Despite a strong link between stressful life events and depression, understanding the mechanisms that underlie the transition from initial stress to a pathological depressive state remains elusive. To date, the majority of preclinical studies have focused on the short-lasting effects of stress, although it is well-recognized that stress can elicit long-lasting and late-appearing effects^15, 14, 32^. In the present study, we characterized the development of affective and cognitive impairments during the days and weeks following social defeat stress to unravel how these core depressive-like symptoms develop. Moreover, as our previous study demonstrated that changes in hippocampal ECM underlie hippocampus-dependent cognitive dysfunction during chronic depression^13^, we here characterized ECM remodeling over time to depict the progression of SDPS-induced hippocampal pathology.

The core depressive symptoms include persistent low mood and anhedonia that reflect abnormal emotion regulation and reward processing^33^. As an example, aberrant social interaction and avoidance of social contact are common in MDD patients^34^. Furthermore, the lack of social interaction can worsen the illness by augmenting stress responses, thereby perpetuating a vicious disease cycle^35^. In preclinical models, social interaction is oftentimes measured as the approach towards a social target, which is a naturally rewarding activity^36^. Here, we found that social deficits develop progressively during the weeks following social defeat stress. Social avoidance behavior early after social defeat stress is generally employed to identify stress-susceptible individuals in mice^37, 38^. In discordance with this, we here show that rats display intact social behavior 24 h after social defeat, which is followed by gradually deteriorating social interaction. Short-term isolation increases the likelihood of social exploration in male rats^39^ and hence this species-specific behavior might have masked the initial deficits in social interaction. On the other hand, the delayed emergence of affective deficits could suggest a gradual progression of the (mal)adaptation underlying emotional regulation in depression. As such, stress-induced early changes might not be accurate in predicting the emergence of enduring affective vulnerability. In support of this, we recently showed that acute effects of defeat stress on social behavior are poor predictors of depression vulnerability^27^. In fact, and in accordance with the present study, our previous data suggest that diminished social interaction displayed 5 weeks post-social defeat most faithfully predicts SDPS susceptibility and depression proneness. These findings indicate that defeat-induced affective deficits develop gradually following social stress and are likely facilitated by prolonged social deprivation^40^ in rats.

Although MDD is thought of as a primarily affective disorder, it is increasingly acknowledged that cognitive dysfunction is common among MDD patients and contributes profoundly to the disease burden^41, 42^. While affective dysfunction and cognitive disturbance co-occur during depressive episodes, these symptoms can also be experienced independently of one another. Cognitive symptoms often persist even in remitted patients and can hinder functional recovery of patients^43^. Moreover, these residual symptoms can serve as a risk factor for relapse or potentially even increase susceptibility for a first depressive episode^44^. Additionally, recent studies suggest that cognitive performance of patients during premedication can predict antidepressant treatment response^45^. Thus, in addition to alleviating mood-affecting symptoms, it is critical to understand and relieve the cognitive deficits in depression, which can sustain disability in patients.

In the current study, spatial memory deficits were present early after the last social defeat encounter. In accordance, several studies have described stress-induced deterioration of hippocampal function shortly after stress^9, 46^. This early phase, i.e. days after stress, is accompanied with substantial structural remodeling of neurons and aberrant hippocampal plasticity, effects that are largely mediated via glucocorticoid signaling^47, 48, 49^. Similar to several other stress paradigms, we have shown that 24 h after social defeat glucocorticoid levels are increased^13^. It is then plausible that the early ECM changes we observed here act additionally, or synergistically, to glucocorticoid-mediated effects that are associated with hippocampal dysfunction and memory deficits^50, 51^.

Strikingly, after these early stress effects subsided, we here showed a period of apparent intact hippocampal function that lasted several weeks before subsequent deterioration. The reversibility of the memory deficit that coincides with recent stress suggests that the late appearing effects evolve gradually and might be mediated by independent neurobiological substrates. In support, early stress effects like increased glucocorticoid response, as well as those on heart rate, body temperature, activity^52^, and water- and food intake, dampen over time and are no longer present during the sustained depressive-like state^13^. Together, these findings highlight a dichotomy in the development of affective and cognitive depression-related symptoms, and, importantly, dissociate more acute stress responses from the late-appearing pervasive deficits that characterize the enduring depressive-like state (**Fig. 5**). This divergent developmental course of affective and cognitive deficits in response to stress likely reflects dichotomous pathophysiological mechanisms underlying these endophenotypes of depression. Accordingly, these two behavioral domains affected in depression may require distinct therapeutic approaches in order to reach full recovery.

**Figure 5.**
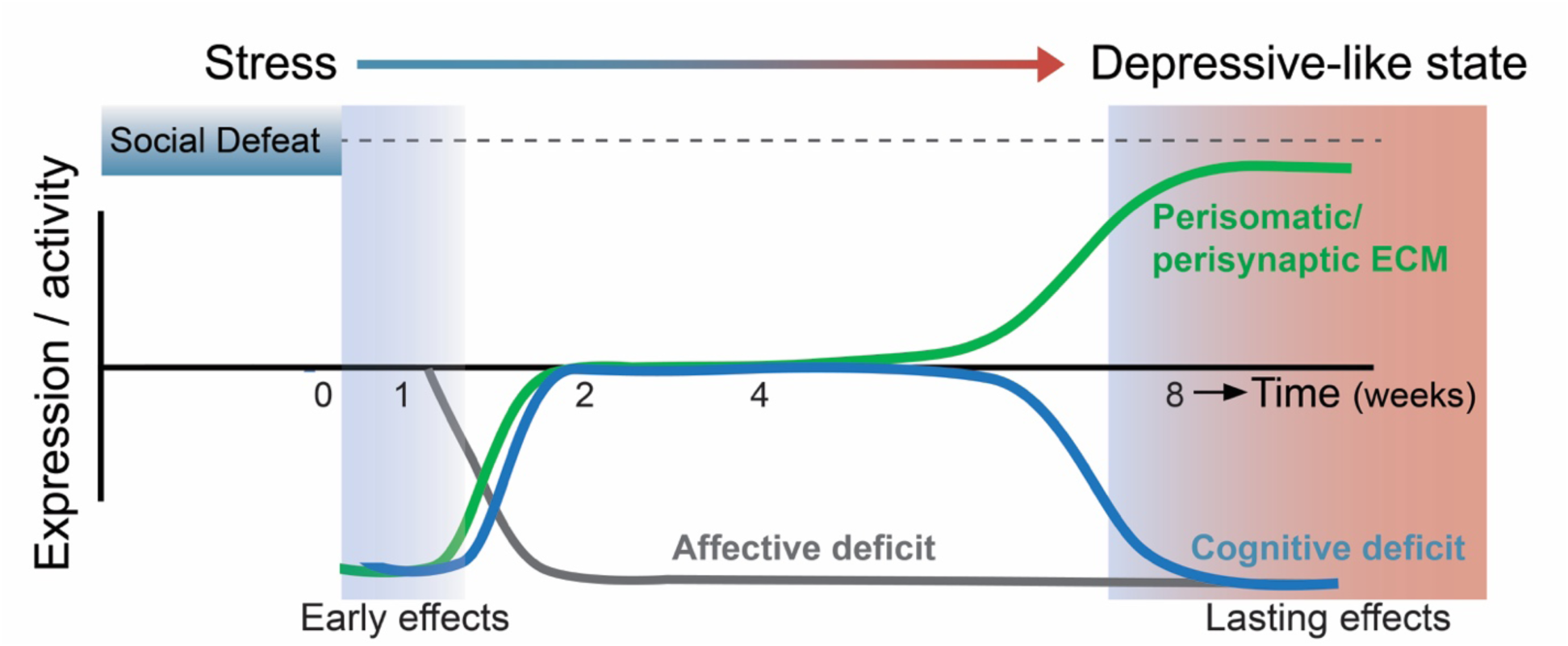
Schematic of the main conclusion: ECM imbalance coincides with cognitive deficits. Overview of the social-stress-induced behavioral impairments and changes in hippocampus ECM during the development from initial stress towards establishing a depressive-like state. Affective (grey) and cognitive (blue) deficits are not temporally associated. Yet, dysregulation of perisomatic (PNN) and perisynaptic ECM (green) coincides with moments of hippocampal dysfunction in terms of spatial memory in the rat SDPS model of depression.

We recently identified changes in hippocampal ECM to underlie hippocampus-dependent memory deficits at the sustained depressive-like state^13^. In the present study, we profiled ECM changes in parallel to the behavioral characterization, in order to understand how aberrant ECM evolves in response to stress. Here, we found social defeat stress to induce an immediate decrease in the expression of synaptic ECM proteins and in the density of perineuronal nets (PNN^+^/PV^+^-cells), without affecting the number of PV^+^-cells. Chondroitin sulfate proteoglycans (CSPGs) are among the most abundant ECM components in the brain, forming mesh-like structures around synapses, and they can condense into PNNs enwrapping predominantly PV^+^-interneurons^53, 54^. Furthermore, the glycoprotein TnR links CSPGs together and Hapln1 attaches CSPGs to the hyaluronan backbone, thereby providing stability to the molecular matrix^55^. Damage to the ECM can destabilize connectivity between neurons and glia, disrupt neural activity and affect neural plasticity^56^; all mechanisms that are critical for learning and memory. For example, genetic deletion of TnR results in changes in perisomatic inhibition and excitability of pyramidal cells that are linked to altered long-term potentiation, the cellular correlate of learning and memory^57^. In addition, TnR ablation impairs spatial and motor learning^58^. Similarly, loss of brevican disrupts long-term potentiation^59^, and results in memory deficits^60^. Together, the downregulation in the expression of several synaptic ECM proteins shortly after social defeat may result in ECM composition-dependent aberrant synaptic transmission that could contribute to the observed memory deficits.

Perineuronal nets are found to mainly enwrap PV^+^-interneurons in the hippocampal CA1 subregion^18^. Although the precise mechanisms by which PNNs regulate the activity of PV^+^-interneurons remains elusive, PNNs have been shown to modulate the excitability and high-frequency firing of PV^+^-interneurons^61, 62^. Recently, it was shown that mainly Brevican that is expressed by PV^+^-interneurons controls PV-excitability via the modulation of excitatory inputs these neurons receive, and controls molecular components of synaptic and intrinsic plasticity^60^. Conversely, hippocampus-dependent learning, and intrinsic plasticity affect activity of PV^+^-interneurons and Brevican levels^63, 60^. These reciprocal relations illustrate that an imbalance in PNNs can alter excitatory/inhibitory signaling as well as network plasticity crucial to cognitive processing^64, 65, 66^. Thus, similar to the increased number of PNNs months after social defeat that coincides with reduced inhibitory network activity and cognitive deficits^13^, the decreased number of PNNs found shortly after defeat stress may alter inhibitory activity of PV^+^-interneurons and thereby impair hippocampus-dependent memory.

The observed co-occurrence of ECM regulation in the synaptic fraction and at the level of PNNs suggests that stress induces a widespread ECM remodeling in the hippocampus that is presumably regulated by the enzymes that breakdown ECM, such as matrix metalloproteinases (MMPs)^30^. Although MMP-9 is mostly studied due to the much wider availability of tools, several MMP-s and modulating enzymes (e.g. tissue inhibitor metalloproteinases, TIMPs) have been shown to be involved in memory deficits^67, 68, 69, 70^. Here, we did not observe major changes in MMP-2 or MMP-9 activity during the early phase after social defeat stress, suggesting that a reduction in the pericellular and -synaptic ECM levels could possibly result from a reduced production or release of ECM proteins, rather than increased breakdown of the ECM by MMP-2 or MMP-9. Although measuring enzyme activity is more informative than measuring MMP-protein levels, the assay might not report endogenous enzyme activity because of possible auto-activation within the gel and loss of endogenous TIMP activity during preparation^71^. Moreover, as we analyzed the whole dorsal hippocampus, we cannot exclude the presence of a subregion-specific increase in MMP-activity shortly after stress, as observed previously^72^. Alternatively, other ECM-cleaving enzymes, such as ADAMTS4/5^73, 74^, could be regulated in response to stress and contribute to the ECM breakdown. It is plausible that proteolysis of ECM during and shortly after stress exposure may promote susceptibility to adverse environmental effects by making the extracellular milieu more permissive to synaptic reorganization. Eventually, this could result in the emergence of a pathological circuitry, which in turn could mediate the long-term changes manifested in late-appearing cognitive deficits.

Interestingly, in the present study we found ECM levels, both at the perisynaptic and perisomatic level, to recover after the initial stress-induced downregulation. Thus, this ECM remodeling displayed a similar profile as hippocampus-dependent memory performance, namely, both showing a full recovery after the acute stress phase before deteriorating at a later phase. Previous studies have demonstrated reversibility of stress-induced hippocampal structural changes, such as an increase in the complexity of dendritic arborization of CA3 neurons and cognitive function after a recovery period following stress that is concurrent with improvement of cognitive deficits^75^. This reversal of the early stress effects has been suggested to represent a dynamic process rather than merely a return to the pre-stress state^76^. Indeed, we describe a transient rescue of stress-induced structural and functional changes at the level of hippocampal ECM and memory performance, respectively. We hypothesize that the initial stress-induced ECM down-regulation possibly triggers a cascade of compensatory mechanisms, involving enzymes responsible for ECM breakdown, together with astrocytic and neuronal ECM production. Presumably these compensatory processes aim to counteract ECM-driven changes in PV activity, which is critical to experience-dependent plasticity and learning^63^. Because SDPS animals are deprived of environmental and social enrichment, it is possible that in these conditions <u>c</u>ompensatory systems fail, in turn contributing to the later emerging pathological state, observed months after social stress.

To shed light on the causal relationship of the early- and late-appearing ECM changes in the SDPS model, it would be critical to investigate whether ECM alterations and cognitive decline observed during the chronic phase of the depression-like state could be attenuated by preventing ECM remodeling during initial stress exposure or shortly thereafter. Although the interdependence of the early- and late-appearing ECM regulation remains under investigation, our findings suggest a critical involvement of ECM remodeling in the progression of stress-induced hippocampal dysfunction as both phases coincided with a corresponding memory deficit (**Fig. 5)**. In support, those time-points for which ECM regulation was absent coincided with intact hippocampal function. Furthermore, as we found MMP-2 activity downregulated during the sustained depressive-like state, this suggests that attenuated MMP-2-mediated ECM breakdown facilitates the build-up of the ECM, thereby promoting the emergence of a state of limited plasticity. In relation to the cognitive phenotype we observe at the long-term, it is of interest that MMP-2 and MMP-9 activity are downregulated in serum of patients with recurrent depression, where higher MMP-levels were correlated with better cognitive performance^77^. Taken together, our findings suggest that ECM remodeling serves as a substrate in mediating both the short- and long-term effects of social stress, and that normal ECM levels are essential for optimal memory processing.

## Conclusion

Collectively, our results show that while SDPS elicits enduring social avoidance and cognitive impairment, these phenotypes progress differently, revealing a temporal dichotomy between affective behavior and cognitive function as often observed in MDD patients^78^. Moreover, our results reveal that SDPS induced a biphasic regulation of cognitive function, highlighting the dissociation of the sustained depressive-like state from the acute effects of stress. Furthermore, as we show that hippocampal ECM remodeling accompanies both early and long-lasting SDPS effects, targeting ECM levels may provide novel therapeutic opportunities against depression and other stress-related neuropsychiatric disorders.

## Acknowledgementss

DR and ABS received support from NBSIK PharmaPhenomics grant LSH framework FES0908; MKK, DR, and SS were supported by an NWO VICI grant (ALW-Vici 016.150.673 / 865.14.002); DR was supported by the EU Research and Innovation Framework Programme Horizon 2020: Marie Skłodowska Curie Actions, Individual Fellowship (MSCA-IF-EF-ST; 793106).

## Author contributions

MKK, DR, ABS, SS designed the experiments

DR, MKK executed behavioral experiments

DR analyzed behavioral data

YvM performed all decapitations for molecular analyses

MKK, DR executed and analyzed molecular data

MKK, SS made the figures

MKK, DR, SS wrote the manuscript

ABS commented during the study and on the manuscript

