## Supplementary material for "From stress to depression: Development of extracellular matrix-dependent cognitive impairment following social stress"

### Contains:

Full method section
Supplemental Figure 1: Correlation of affective and cognitive phenotypes
Supplemental Figure 2: Whole immunoblot compilation 72 h post-defeat
Supplemental Figure 3: Whole immunoblot compilation week 3 post-defeat
Supplemental Figure 4: Whole gel gelatinase assay 72 h and week 8 post-defeat

Supplemental Table 1: Overview statistics of all main figures.
<u>Note</u>: Supplemental Table 2: Overview data of all main figures. This is provided separately as excel file.

### Full method section

Animals and the social defeat-induced persistent stress paradigm

All experiments were approved by the Vrije University Amsterdam Animal Users Committee and in accordance with the relevant guidelines and regulations. Wistar male rats, 7 weeks old and weighing <200 g upon arrival (Envigo, The Netherlands) were pair-housed and allowed to habituate to the facility for at least 2 weeks before the start of the experiments. Long-Evans male rats (Charles River, UK) were used as residents for this resident-intruder paradigm. The residents were pair-housed with sterilized females >1 week before the start of the experiment. Females were removed from the resident's cage prior to the defeat sessions. The SDPS rats underwent 15-minute social defeat sessions daily. During the defeat, the rats were first placed inside the defeat apparatus, while a transparent perforated plexiglass partition wall separated the intruder from the resident, permitting sensory exchange but preventing all physical contact for 5 minutes. The partition wall was then removed and a 5-minute fight phase started. Following the defeat, the rats were separated again and sensory exchange was allowed for another 5-minutes. Social defeat was repeated for five consecutive days and each day a new resident was used. From the first defeat session onwards, the SDPS rats were single-housed and kept in social isolation until the end of the experiments.

Behavioral testing

All animals were habituated to the testing arenas (79 x 57 x 42 cm, plastic) before they underwent any behavioral testing. During the habituation phase, the rats were transported to the video-recording room and let to freely explore an empty testing arena for 10 minutes. The habituation was conducted for 3 times. All behavioral testing was performed during the dark phase under a dim red light.

*Social Approach Avoidance – SAA test:* The SAA test consisted of 3 phases: habituation, sampling and testing. During habituation, the animals were allowed to explore the empty arena for 5 minutes. This was followed by sampling phase when two empty interaction boxes (23 x 11 x 34 cm, perforated, transparent) were introduced to the arena and the animals were allowed to explore the empty boxes for 5 minutes. After the sampling phase, the rats were placed back to their home-cage, and an unfamiliar Long-Evans rat was placed in one of the boxes. Immediately thereafter, the animals were introduced back to the arena and were allowed to explore the arena again for 5 minutes. The interaction ratio was calculated as time spent near the social target vs. time spent near the empty box: social target zone / (social target zone + empty box zone) during the first minute of the test.

*Object Place Recognition – OPR test*: The OPR test consisted of 3 phases: habituation, sampling and testing. During the habituation, the animals were allowed to explore an empty arena for 5 minutes. This was followed by sampling phase upon which two identical objects (8 x 8 x 35 cm, metal, cylinders or cubes) were introduced to the testing arena. After the sampling phase, the rats were placed back to their home-cage. Two objects were replaced with a set of two identical objects but one object was relocated to a different corner within the arena. After a 15-minute interval, the rats were introduced back to the arena and exploration of the objects was analyzed for the first 1 minute. The discrimination index was calculated by measuring the time spent exploring the relocated object compared to the stable object: relocated object / (relocated object + stable object).

Immunohistochemistry

Following transcardial perfusion with and overnight post-fixation in ice-cold 4% PFA in PBS, brains were transferred to 30% sucrose in PBS in 4 °C until sectioned. Free-floating cryostat sections (35 µm) were collected from the dorsal hippocampus (AP -2.40 to -4.56) and stored in PBS+0.02% $NaN_3$ until further use. Sections were washed 3 times for 10 minutes in PBS, followed by a blocking for 1 h at RT in blocking solution (2.5% BSA, 0.1% Triton, 5% goat serum in PBS). After blocking, the sections were incubated with primary antibodies (mouse anti-chondroitin sulfate proteoglycan 1:1000, cat-301 MAB5284; rabbit anti-parvalbumin 1:1000, Swant #235) overnight at 4 °C. This was followed by washing in PBS 4 times 10 minutes at RT, and incubation with fluorescent-conjugated secondary antibodies (anti-mouse-Alexa-488 1:400, Invitrogen A11001; anti-rabbit-Alexa-568 1:400, Invitrogen A11011) for 2 h at RT. Thereafter, the sections were washed 4 times 10 min with PBS at RT. Sections were mounted onto microscope slides using PBS+0.02% gelatin and coverslipped with DAPI-containing mounting medium (H-1500 Vector Laboratories).

Images were acquired on a fluorescent microscope (Leica DM5000). The Fiji program, using automated threshold and particle analysis, was used to detect the number of $PNN^+$ and $PV^+$ cells and their double immunoreactivity. False-positive cells were excluded manually during the analysis. During the image acquisition and cell quantification, the researcher was blind to the experimental groups.

Tissue preparation – synaptosomes

Following decapitation, the brain was removed, and the dorsal hippocampus was immediately dissected on ice[1], and frozen on dry ice. Samples were homogenized in ice-cold 0.32 M sucrose (5% of homogenate was collected as total cell lysate) and then centrifuged at 1000x *g* for 10 minutes. The supernatant was loaded on top of a sucrose gradient consisting of 0.85 and 1.2 M sucrose. After centrifugation at 100,000x *g* for 2 h, the synaptosome fraction at the interface of 0.85/1.2 M sucrose was collected and then lysed in hypotonic solution.

**Statistics**

All data were analyzed using IBM SPSS Statistics 24. For group comparisons, two-tailed Student's t-tests (with or without correction for unequal variation) were applied for normally distributed data and Mann-Whitney U-tests otherwise. Data were checked for normality using the Saphiro-Wilk test.

All group data are depicted as mean±SEM, with individual data on top. Statistical significance level was set for *P*-values<0.05, trend for 0.05<*P*-value<0.10. Details of all statistical testing can be found in Supplementary Table S1, and individual data points can be found in Supplementary Table S2.

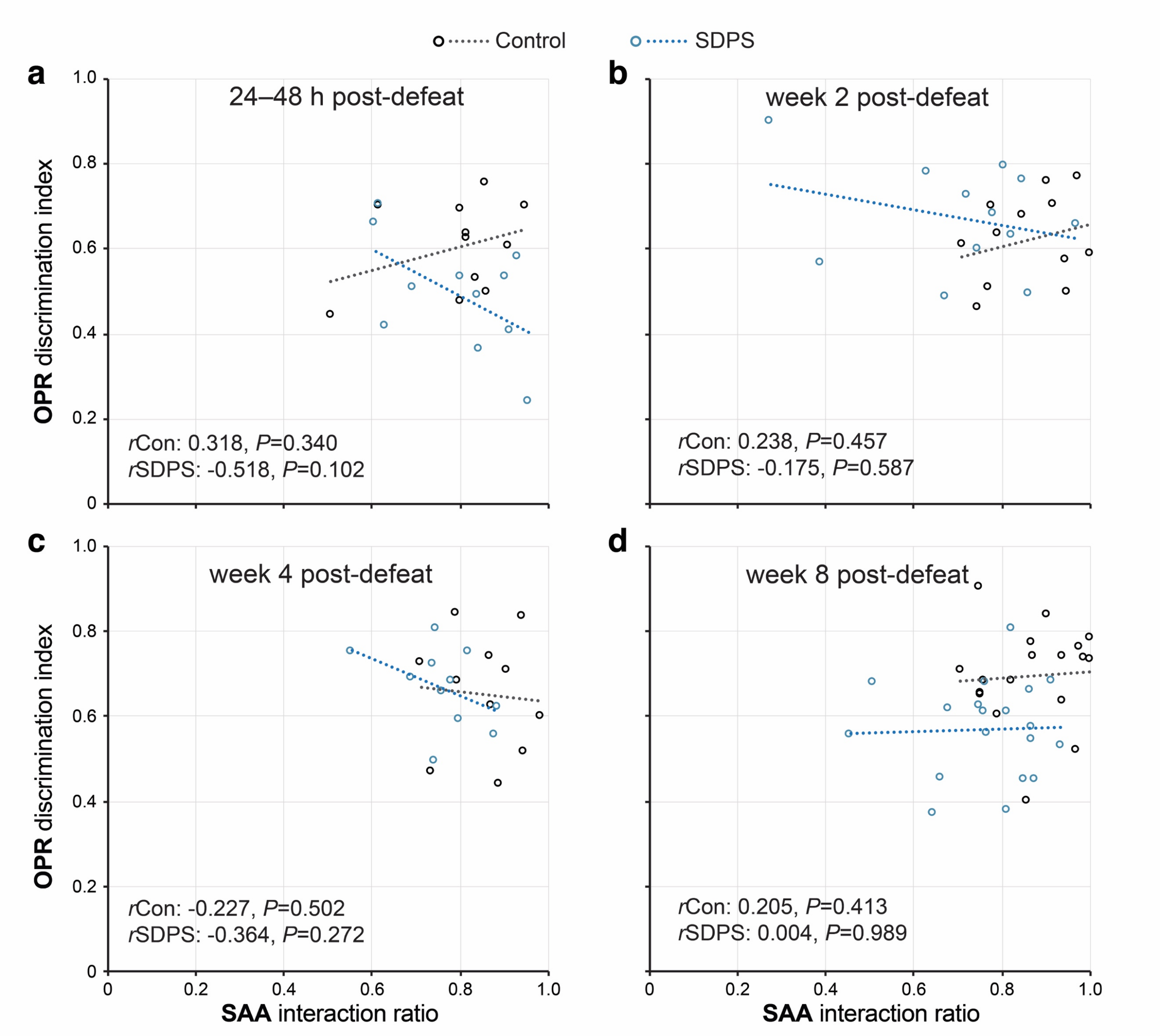


**Supplemental Figure 1: Correlation of affective and cognitive phenotypes.** SAA interaction ratio and OPR discrimination index were correlated per animal batch undergoing SAA and OPR behavior, either shortly after defeat (**a**), at week 2 (**b**), week 4 (**c**) and week 8 (**d**) post-defeat. Spearman correlations (*r*) and *P*-values are indicated per group (Con, SDPS), as well as the trendline (dotted line). In addition, Pearson correlation over the entire group per time point indicated the absence of any significant correlations between affective and cognitive parameters ($r_{24\text{-}48\ h}$: -0.146 *P*=0.517; $r_{2\ w}$:-0.212 *P*=0.319; $r_{4\ w}$: -0.223 *P*=0.318; $r_{8\ w}$:0.233 *P*=0.165; see Supplemental Table 1).

### Supplemental material

Koskinen et al. – From stress to depression: Development of extracellular matrix-dependent cognitive impairment following social stress

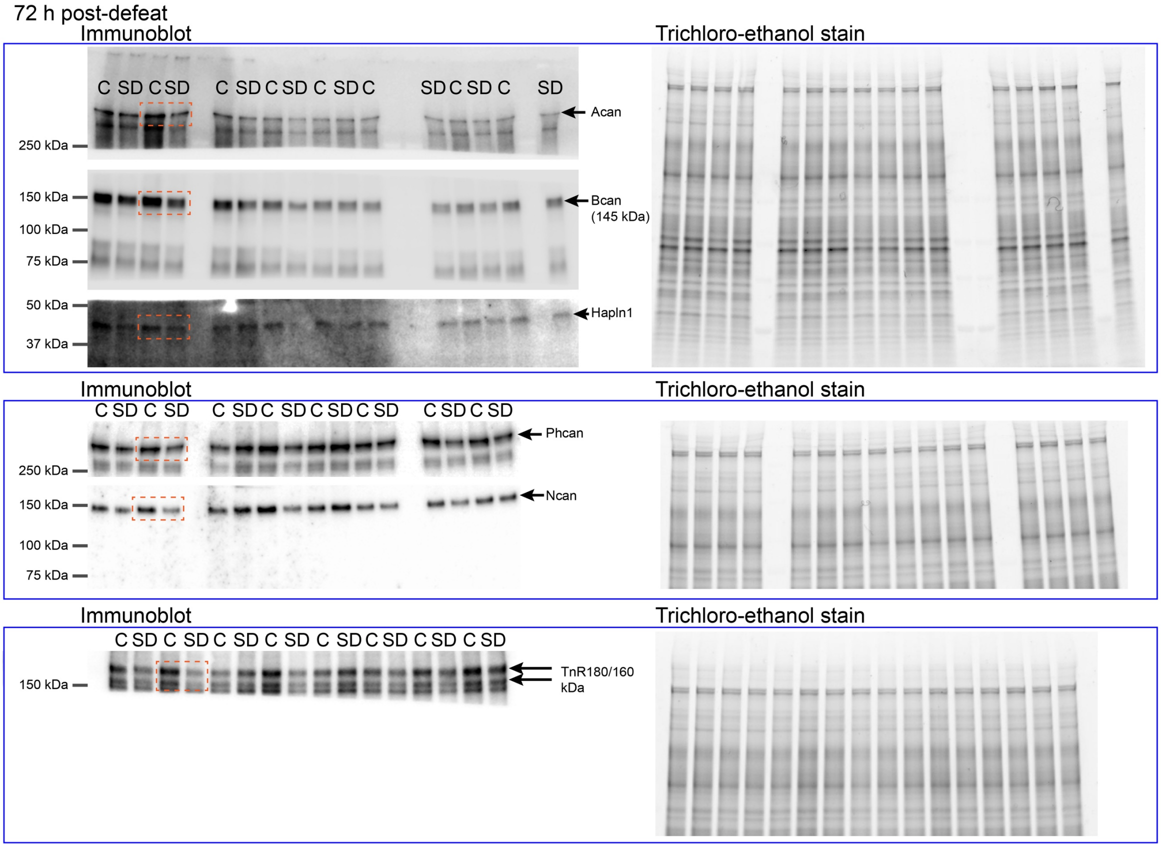


**Supplemental Figure 2. Whole immunoblot compilation 72 h post defeat.** Whole immunoblots (left) and trichloro-ethanol stained gels (right) are shown from which sections (orange dashed rectangle) are included in Figure 3 for the 72 h post-defeat time-point (C=control animal; SD=SDPS animal). Molecular weights are indicated.

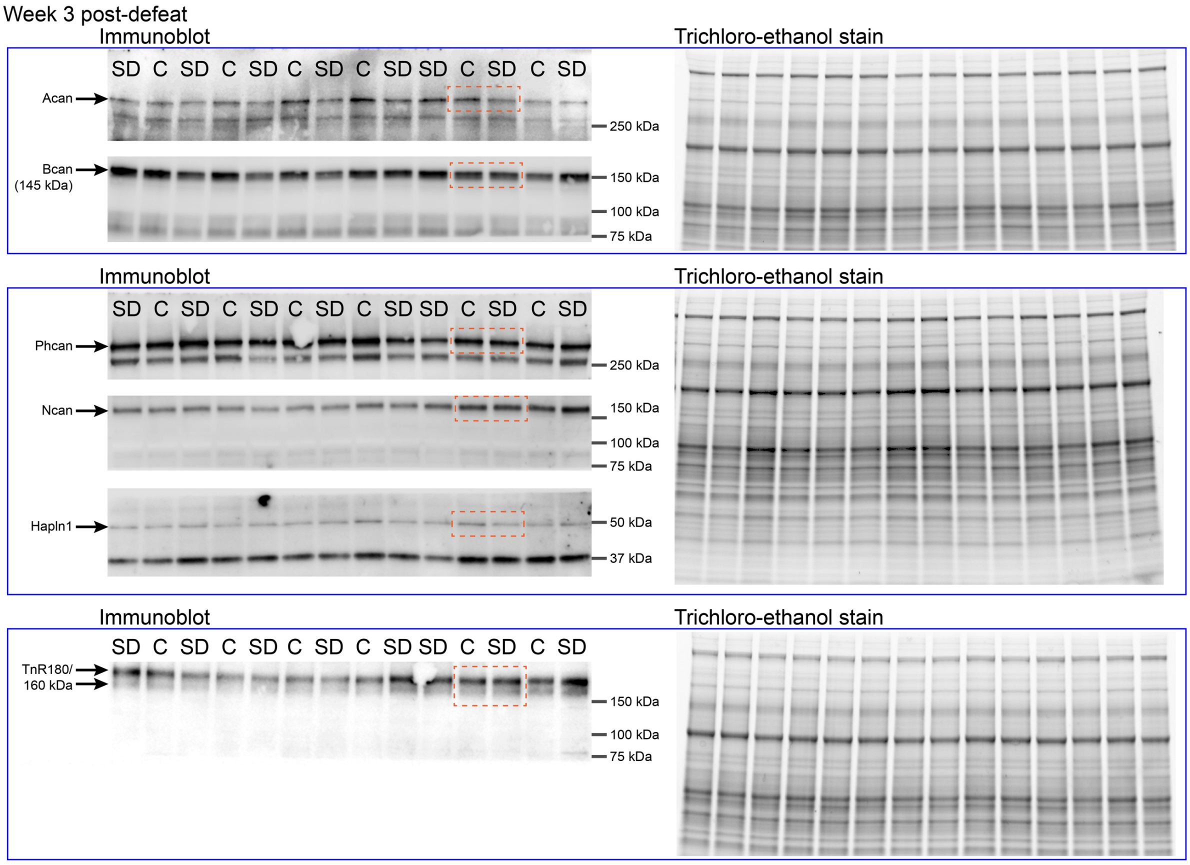


**Supplemental Figure 3. Whole immunoblot compilation week 3 post defeat.** Whole immunoblots (left) and trichloro-ethanol stained gels (right) are shown from which sections (orange dashed rectangle) are included in Figure 3 for the 3 week post-defeat time-point (C=control animal; SD=SDPS animal). Molecular weights are indicated.

**Supplemental material**
Koskinen et al. – From stress to depression: Development of extracellular matrix-dependent cognitive impairment following social stress

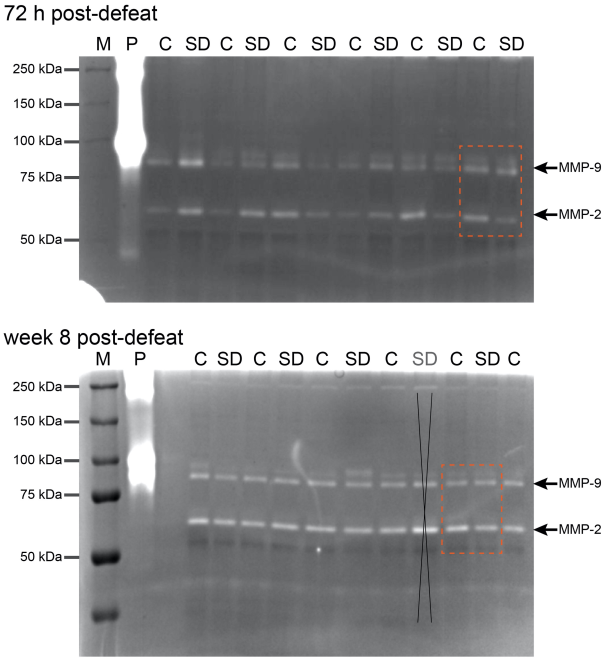


**Supplemental Figure 4. Whole gel gelatinase assay 72 h and week 8 post-defeat.** Whole gels for in gel zymography (72 h, up; week 8, down) are shown from which sections (orange dashed rectangle) are included in in Figure 4. The mature form of MMP-2 and MMP-9 (white bands), as well as molecular weights are indicated. Sample normalization for loading differences was performed on the Coomassie-stained bands. M, marker; P, positive control of human MMP-9, C=control animal; SD=SDPS animal. Note that 1 SDPS sample (crossed, week 8) was considered an outlier at the protein level and for MMP activity.

**Supplemental material**
Koskinen et al. – From stress to depression: Development of extracellular matrix-dependent cognitive impairment following social stress

**Supplemental Table 1. Overview statistics all figures.** Shown are the results and details of the 2-tailed statistical analyses (type of test, n-number, t-value, Df, *P*-value) for the data shown in Figure 1 (behavior) and Supplemental Figure 1 (correlation of behavior), Figure 2 (PNN staining), Figure 3 (Immunoblot), and Figure 4 (MMP gelatinase activity) in SDPS *vs.* Control animals for the indicated time points. Significance (*P*<0.050) is indicated in bold, trend (*P*<0.100) in underlined; tests with unequal variance are in italics.

| ***Figure*** | ***Statistical test*** *($n_{Con}$, $n_{SDPS}$)* | ***Statistics*** *(t-value, Df)* | ***P-value*** |
|---|---|---|---|
| 1B | **Mann-Whitney U test** | | |
| | CON vs. SDPS 24 h (11, 11) | | *P* = 0.935 |
| | CON *vs.* SDPS week 8 (18, 19) | | ***P* = 0.026** |
| | **Unpaired t-test** | | |
| | CON *vs.* SDPS week 2 (12, 12) | t(22) = 2.352 | ***P* = 0.028** |
| | CON *vs.* SDPS week 4 (11, 11) | t(20) = 2.456 | ***P* = 0.023** |
| | CON *vs.* fictive 24 h | (t20) = 5.484 | **$P < 10^{-4}$** |
| | CON *vs.* fictive week 2 | t(22) = 8.888 | **$P < 10^{-4}$** |
| | CON *vs.* fictive week 4 | t(20) = 9.457 | **$P = 10^{-4}$** |
| | CON *vs.* fictive week 8 | t(34) = 11.104 | **$P < 10^{-4}$** |
| | SDPS *vs.* fictive 24 h | t(20) = 5.162 | **$P < 10^{-4}$** |
| | SDPS *vs.* fictive week 2 | t(22) = 2.575 | ***P* = 0.017** |
| | SDPS *vs.* fictive week 4 | t(20) = 6.733 | **$P < 10^{-4}$** |
| | SDPS *vs.* fictive week 8 | t(36) = 6.382 | **$P < 10^{-4}$** |
| 1C | **Unpaired t-test** | | |
| | CON *vs.* SDPS 48 h (11, 11) | t(20) = 2.189 | ***P* = 0.041** |
| | CON *vs.* SDPS week 2 (12, 12) | t(22) = -1.067 | *P* = 0.298 |
| | CON *vs.* SDPS week 4 (11, 11) | t(20) = -0.271 | *P* = 0.789 |
| | CON *vs.* SDPS week 8 (18, 19) | t(35) = 3.366 | ***P* = 0.002** |
| | CON *vs.* fictive 48 h | t(20) = 2.349 | ***P* = 0.029** |
| | CON *vs.* fictive week 2 | t(22) = 2.944 | ***P* = 0.008** |
| | CON *vs.* fictive week 4 | t(20) = 2.583 | ***P* = 0.018** |
| | CON *vs.* fictive week 8 | t(34) = 5.094 | ***P* < 0.001** |
| | SDPS *vs.* fictive 48 h | t(20) = -0.116 | *P* = 0.908 |
| | SDPS *vs.* fictive week 2 | t(22) = 3.356 | ***P* = 0.003** |
| | SDPS *vs.* fictive week 4 | t(20) = 4.106 | ***P* = 0.001** |
| | SDPS *vs.* fictive week 8 | t(36) = 1.890 | *P* = 0.067 |
| S1A–D | **Pearson correlation SAA & OPR per time point** | | |
| | $SAA_{24\ h}$ *vs* $OPR_{48\ h}$ (22) | $r_{24\text{-}48\ h}$ = -0.146 | *P* = 0.517 |
| | $SAA_{2\ w}$ *vs* $OPR_{2w}$ (24) | $r_{2\ w}$ = -0.212 | *P* = 0.319 |
| | $SAA_{4\ w}$ *vs* $OPR_{4\ w}$ (22) | $r_{4\ w}$ = -0.223 | *P* = 0.318 |
| | $SAA_{8\ w}$ *vs* $OPR_{8\ w}$ (37) | $r_{8\ w}$ = 0.233 | *P* = 0.165 |
| | **Spearman correlation SAA & OPR per group per time point** | | |
| | **Control** | | |
| | $SAA_{24\ h}$ *vs* $OPR_{48\ h}$ (11) | $r_{24\text{-}48\ h}$ = 0.318 | *P* = 0.340 |
| | $SAA_{2\ w}$ *vs* $OPR_{2w}$ (12) | $r_{2\ w}$ = 0.238 | *P* = 0.457 |
| | $SAA_{4\ w}$ *vs* $OPR_{4\ w}$ (11) | $r_{4\ w}$ = -0.227 | *P* = 0.502 |
| | $SAA_{8\ w}$ *vs* $OPR_{8\ w}$ (18) | $r_{8\ w}$ = 0.205 | *P* = 0.413 |
| | **SDPS** | | |
| | $SAA_{24\ h}$ *vs* $OPR_{48\ h}$ (11) | $r_{24\text{-}48\ h}$ = -0.518 | *P* = 0.102 |
| | $SAA_{2\ w}$ *vs* $OPR_{2w}$ (12) | $r_{2\ w}$ = -0.175 | *P* = 0.587 |
| | $SAA_{4\ w}$ *vs* $OPR_{4\ w}$ (11) | $r_{4\ w}$ = -0.364 | *P* = 0.272 |
| | $SAA_{8\ w}$ *vs* $OPR_{8\ w}$ (19) | $r_{8\ w}$ = 0.004 | *P* = 0.989 |
| 2B | **Unpaired t-test** | | |
| | CON *vs.* SDPS 72 h (5, 6) | t(9) = 2.569 | ***P* = 0.030** |
| | CON *vs.* SDPS week 2 (4, 4) | t(6) = 0.077 | *P* = 0.941 |
| | CON *vs.* SDPS week 4 (3, 4) | t(5) = -1.490 | *P* = 0.196 |
| | CON *vs.* SDPS week 8 (4, 4) | t(6) = -3.087 | ***P* = 0.021** |
| 2D | **Unpaired t-test** | | |
| | CON *vs.* SDPS 72h | t(5) = -0.030 | *P* = 0.977 |
| | CON *vs.* SDPS week 3 | t(5) = -1.673 | *P* = 0.155 |
| | CON *vs.* SDPS week 5 | t(5) = -1.940 | *P* = 0.110 |
| | CON *vs.* SDPS week 9 | *t(3.850) = -1.105* | *P = 0.333* |

| | | | |
|---|---|---|---|
| 3B,C | *Log$_2$ transformed data* | | |
| | **72 h** | | |
| | **Unpaired t-test** | | |
| | CON *vs*. SDPS Acan (8, 7) | t(13) = 0.779 | *P* = 0.450 |
| | CON *vs*. SDPS Bcan (8, 7) | t(13) = 2.300 | ***P* = 0.039** |
| | CON *vs*. SDPS Ncan (8, 7) | t(13) = 2.453 | ***P* = 0.029** |
| | CON *vs*. SDPS Pcan (8, 7) | t(13) = 2.610 | ***P* = 0.022** |
| | CON *vs*. SDPS TenR160 (8, 6) | t(13) = 3.213 | ***P* = 0.007** |
| | CON *vs*. SDPS TenR180 (8, 7) | t(13) = 2.197 | ***P* = 0.047** |
| | **Mann-Whitney U test** | | |
| | CON *vs*. SDPS Hapln1 (8, 7) | | ***P* = 0.021** |
| | **Week 2** | | |
| | **Unpaired t-test** | | |
| | CON *vs*. SDPS Acan (6, 8) | t(12) = 1.083 | *P* = 0.300 |
| | CON *vs*. SDPS Bcan (6, 8) | t(12) = 0.009 | *P* = 0.993 |
| | CON *vs*. SDPS Ncan (6, 8) | t(12) = -0.312 | *P* = 0.760 |
| | CON *vs*. SDPS Pcan (6, 8) | t(12) = 0.435 | *P* = 0.671 |
| | CON *vs*. SDPS TenR160 (6, 8) | t(12) = -1.021 | *P* = 0.327 |
| | CON *vs*. SDPS TenR180 (6, 8) | t(12) = -0.495 | *P* = 0.629 |
| | CON *vs*. SDPS Hapln1 | t(12) = 0.710 | *P* = 0.491 |
| | **Week 8*** | | |
| | **Unpaired t-test** | | |
| | CON *vs*. SDPS Acan (4, 5) | t(7) = -1.107 | *P* = 0.305 |
| | CON *vs*. SDPS Bcan (4, 5) | t(7) = -3.582 | ***P* = 0.009** |
| | CON *vs*. SDPS Pcan (4, 5) | t(7) = -3.563 | ***P* = 0.009** |
| | CON *vs*. SDPS TenR160 (4, 4) | t(7) = -1.999 | *P* = 0.093 |
| | CON *vs*. SDPS TenR180 (4, 4) | t(7) = -1.505 | *P* = 0.183 |
| | CON *vs*. SDPS Hapln1 | t(7) = -3.226 | ***P* = 0.015** |
| | **Mann-Whitney U test** | | |
| | CON *vs*. SDPS Ncan (4, 5) | | ***P* = 0.016** |
| 4,B,C | **72 h** | | |
| | **Unpaired t-test** | | |
| | CON *vs*. SDPS MMP-2 (6, 6) | t(10) = -0.513 | *P* = 0.619 |
| | CON *vs*. SDPS MMP-9 (6, 6) | t(10) = -1.09 | *P* = 0.301 |
| | **Week 8** | | |
| | **Unpaired t-test** | | |
| | CON *vs*. SDPS MMP-2 (6, 4) | t(8) = 2.398 | *P* = 0.043 |
| | CON *vs*. SDPS MMP-9 (6, 4) | t(8) = 2.272 | *P* = 0.239 |

* Note, the 9-week immunoblot data (non-log2-transformed) have been published before[2], and are not repeated to adhere to the 3R-principle of animal research.

**Supplemental Table 2. Overview data all main figures.** Shown are all individual data points of Figures 1–4 (*excel file*).
